# Sex Differences in Pubertal Circadian and Ultradian Rhythmic Development Under Naturalistic Conditions

**DOI:** 10.1101/2021.10.15.464452

**Authors:** Azure D. Grant, Linda Wilbrecht, Lance J. Kriegsfeld

## Abstract

Biological rhythms in core body temperature (CBT) provide informative markers of adolescent development under controlled laboratory conditions. However, it is unknown if these markers are preserved under more variable naturalistic conditions, and if CBT may therefore prove useful in a real-world setting. To evaluate this possibility, we examined fecal steroid concentrations and CBT rhythms from pre-adolescence (p26) through early adulthood (p76) in intact male and female rats under natural light and climate at the University of California, Berkeley Field Station. Despite greater environmental variability, CBT markers of pubertal onset and its rhythmic progression were comparable to those previously reported in laboratory conditions in female rats and extend actigraphy-based findings in males. Specifically, sex differences emerged in circadian rhythm (CR) power and temperature amplitude prior to pubertal onset and persisted into early adulthood, with females exhibiting elevated CBT and decreased CR power compared to males. Within-day (ultradian rhythm; UR) patterns also exhibited a pronounced sex difference associated with estrous cyclicity. Pubertal onset, defined by vaginal opening, preputial separation, and sex steroid concentrations, occurred later than previously reported under lab conditions for both sexes. Vaginal opening and increased fecal estradiol concentrations were closely tied to the commencement of 4-day oscillations in CBT and UR power in female rats. By contrast, preputial separation and the first rise in testosterone concentration were not associated with adolescent changes to CBT rhythms in male rats. Together, males and females exhibited unique temporal patterning of CBT and sex steroids across pubertal development, with tractable associations between hormonal concentrations, external development, and temporal structure in females. The preservation of these features outside the laboratory supports CBT as a strong candidate for translational pubertal monitoring under naturalistic conditions in females.

## Introduction

Clinical or self-assessment(Elchuri and Momen, 2020) of pubertal status is typically conducted via observation of external characteristics (e.g., Tanner Scale, developed in the late 1960’s)(Rueda-Quijano et al., 2019; Shirtcliff et al., 2009), menstrual cycle tracking in girls(Fowler et al., 2020), or costly hormone measurement(Klein et al., 2017). Today, relatively inexpensive wearable sensors can capture metrics that are closely influenced by reproductive hormones and metabolism, such as body temperature(Grant et al., 2020; Smarr et al., 2020). These sensors may provide a convenient method to monitor pubertal development in real-world settings. Such sensors, along with a database of normative changes, could provide non-invasive information about pubertal development to teens, families, or clinicians(Wartella et al., 2016). These investigations require years of future study, wider adoption of wearables by preteens and teens, and further development of regulatory standards for wearable companies and clinicians(Campbell-Page and Shaw-Ridley, 2013; Gear, 2014; Grant et al., 2019; Wartella et al., 2016). Although the value of identifying these features for pre-clinical and translational studies is evident, whether environmental variability masks the patterns identified under controlled laboratory conditions requires empirical investigation.

Biological rhythms in core body temperature (CBT) change markedly across adolescence in rodents, enabling unobtrusive monitoring of this trajectory in a laboratory setting(Grant et al., 2021; Hagenauer et al., 2011; Zuloaga et al., 2009). These rhythms are coupled across physiological systems(Goh et al., 2019; Grant et al., 2020, 2018; Mohawk et al., 2012) and at multiple timescales, including within-a-day (ultradian rhythms; URs)(Bourguignon, 1988) and daily (circadian rhythms; CRs)(Garcia et al., 2001; MacKinnon et al., 1978) in both sexes, and multi-day ovulatory cycles in females (ovulatory rhythms; ORs)(Vidal, 2017). Rhythmicity serves numerous functions, including coordination of reproductive development(Albertsson-Wikland et al., 1997; Ankarberg and Norjavaara, 1999; Hagenauer et al., 2011; Norjavaara et al., 1996) and synchronization of internal systems to variation in the environment(Daan and Slopsema, 1978; Hoogenboom et al., 1984; Lewis and Curtis, 2016). Features of biological rhythmicity can provide clinically-relevant diagnostic information(Akin and Elstein, 1975; Bhavani et al., 2019; Grant et al., 2020; Smarr et al., 2020). We recently applied this approach to monitor female adolescent development in rats under controlled laboratory conditions(Grant et al., 2021). This strategy revealed rhythmic features CBT that can be used to track adolescent development, with CR power and CBT amplitude rising from early to mid-adolescence and stabilizing by early adulthood. Such outputs were coordinated with changes in reproductive hormones, consistent with the established temperature-modulating effects of estrogen(Williams et al., 2010) and progesterone(Buxton and Atkinson, 1948).

However, exposure to the greater spectro-temporal variability of natural light, temperature, humidity, and enriched sensory complexity of a naturalistic environment (Joyce et al., 2020; Stothard et al., 2017) may add ‘noise’ to these features, and affect pubertal timing and tempo. Although a great deal of research has focused on extreme environments (e.g., polar(Steiger et al., 2013)), temperate environments may reveal differences from laboratory-derived features. Mice and rats exposed to longer or variable day lengths, for example, exhibit delayed external markers of pubertal onset(Lafaille et al., 2015), more variable activity rhythms(Kim and Harrington, 2008; Meijer et al., 2010), and have altered weight gain trajectories(Brown-Douglas et al., 2004). In contrast, male Siberian hamsters (*Phodopus sungorus*) advance puberty in long day lengths(Park et al., 2003) to maximize reproductive success prior to winter. These changes suggest species-specific decoupling of maturation mechanisms that are coordinated under laboratory conditions and that may also decouple temperature features from sexual maturation (Silva and Domínguez, 2020). Additionally, animals raised in naturalistic environments exhibit elevated steroid hormone concentrations(Woodruff et al., 2013, 2010), suggesting that the hormonal milieu influencing the adolescent trajectory may alter temperature rhythms relative to laboratory-based studies.

To assess the potential impact of these factors on CBT rhythmicity during adolescence, we examined reproductive hormones and CBT patterns in a naturalistic setting. As humans face a complex environment of combined artificial and natural stimuli, we chose to investigate animals housed at the Field Station (FS) in Berkeley, CA, which is an intermediate between laboratory and field conditions. This environment provides shelters open to natural changes in light, humidity, and temperature, as well as a social partner and standard laboratory housing and food. We hypothesized that the FS environment would result in higher and more variable sex steroid concentrations(Woodruff et al., 2013, 2010) and pubertal timing onset compared to previous reports in the laboratory environment(Grant et al., 2021). We also speculated that these changes would be mirrored in CR and UR patterns and temperature amplitude. Finally, we anticipated that reported features of adolescence would occur in males as well as females, with the exception of the emergence of patterns associated with the ovulatory cycle, and that males may exhibit the sex difference of elevated ultradian power and decreased temperature, as previously reported (Zuloaga et al., 2009), compared to females.

## Materials and Methods

### Animals

Male and female Wistar rat breeders were purchased at 250 g and 300 g, respectively, from Charles River (Charles River, Wilmington, MA). Animals were bred at the FS and weaned at postnatal day 21 (p21), with a maximum of one pair of pups (one experimental and one partner pup) in each experimental group, per litter. Weanlings were housed in same-sex pairs to minimize social isolation stress known to affect pubertal development(Bakshi and Geyer, 1999; Boggiano et al., 2008) in standard translucent propylene (96 x 54 x 40 cm) rodent cages, and provided *ad libitum* access to food and water, wood chips for floor cover, bedding material, and chew toys for the duration of the study. Animals were gently handled daily before weighing to minimize stress. To prevent mixing of feces collected, cage mates were separated by a flexible stainless-steel lattice that permitted aural, scent, and touch interaction between siblings. A total of 16 animals were included in the study (n=8 per sex), with 16 same-sex individuals as social, littermate partners. The experiment was conducted in rooms with natural light (light intensity during the mean photo- and scotophases were 677 ± 254 and 2.65 ± 0.40 lux, respectively), outdoor ambient temperatures averaging 22.6 ± 0.34° C , and air circulation from August 9^th^ to September 29^th^, 2019, at the Field Station at the University of California, Berkeley. All procedures were approved by the Institutional Animal Care and Use Committee of the University of California, Berkeley and conformed to the principles in the Guide for the Care and Use of Laboratory Animals, 8^th^ ed.

### Core Body Temperature Data Collection

Data were gathered with G2 E-Mitter implants that chronically record CBT (Starr Life Sciences Co., Oakmont, PA). At weaning, G2 E-Mitters were implanted in the intraperitoneal cavity under isoflurane anesthesia, with analgesia achieved by subcutaneous injections of 0.03 mg/kg buprenorphine (Hospira, Lake Forest, IL) in saline. Buprenorphine was administered every 12 h for 2 days following surgery. E-Mitters were sutured to the ventral muscle wall to maintain consistent core temperature measurements. Recordings began immediately, but data collected for the first 5 days post-surgery were not included in analyses to allow for post-surgical recovery. Recordings were continuous and stored in 1-min bins.

### Fecal Sample Collection

Fecal E2 (fE2) concentrations in females, and fecal testosterone (fT) concentrations in males, were assessed across puberty from feces generated over 24 h periods. Feces provide a more representative sample of average daily hormone concentrations than do blood samples(Auer et al., 2020; Harper and Austad, 2000; Millspaugh and Washburn, 2003; Touma et al., 2004; Woodruff et al., 2010) and are non-invasively generated, thereby reducing stress associated with high-frequency, longitudinal blood collection. Samples were collected in small, airtight bags in the early mornings from p25 to p37 (pre puberty and first estrous cycle), p45 to p51 (mid puberty), and p55 to p65 (late puberty to early adulthood) in females, and every 3 days in males from p25 to p74. Samples soiled with urine were discarded and all other boli generated over each 24-h segment were combined. Samples were stored at - 20° C within 1 h of collection until preprocessing for the ELISA assay. Sample collection took ~ 1 min per animal. One female’s samples were frequently soiled with urine and were therefore not included in analyses of 12 out of 24 of collected timepoints.

Samples were processed according to manufacturer’s instructions (Arbor Assays, Ann Arbor, MI.). Briefly, samples were placed in a tin weigh boat and heated at 65°C for 90 minutes, until completely dry. Dry samples were ground to a fine powder in a coffee grinder, which was wiped down with ethanol and dried between samples to avoid cross contamination. Powder was weighed into 0.2 mg aliquots and added to 2 mL test tubes. For hormone extraction, 1.8mL of 100% ethanol was added to each test tube, and tubes were shaken vigorously for 30 minutes. Tubes were then centrifuged at 5,000 RPM for 15 minutes at 4°C. Supernatant was moved to a new tube and evaporated under 65°C until dry (~ 90 minutes). Sample residue was reconstituted in 100μL of 100% ethanol. 25μL of this solution was diluted for use in the assay and remaining sample was diluted and stored.

### Hormone Assessment

A commercially available fE2 enzyme-linked immunosorbent assay (ELISA) kit was used to quantify E2 in fecal samples (Arbor Assays, Ann Arbor, MI). These assays have been previously published in species ranging from rats and mice(Asimes et al., 2018; Auer et al., 2020; Kalliokoski et al., 2015; Lv et al., 2020; Mathew et al., 2017; Steadman, 2019; Steadman et al., 2019), to wolves(Franklin et al., 2020), to humans(Righetti et al., 2020). ELISAs were conducted according to the manufacturer’s instructions. To ensure each sample contained ≤ 5% alcohol, 25μL of concentrate were vortexed in 475μL Assay Buffer. All samples were run in duplicate, and an inter-assay control was run with each plate. Sensitivity for the estradiol assay was 39.6 pg/mL and the limit of detection was 26.5 pg/mL. Sensitivity for the testosterone assay was 9.92 pg/mL and the limit of detection was 30.6 pg/mL. Fecal testosterone intra-assay coefficient of variation (C.V.) was 9.35% and inter-assay C.V. was 10.5%. Fecal estradiol intra-assay CV was 5.0% and inter-assay CV was 5.54%.

### Data Availability and Analysis

All code and data used in this paper are available at A.G.’s and L.K.’s Github(azuredominique, 2021; Kriegsfeld-Lab, 2021). Code was written in MATLAB 2020b and 2021a with Wavelet Transform (WT) code modified from the Jlab toolbox and from Dr. Tanya Leise(Leise, 2015, 2013). Briefly, data were imported to MATLAB at 1-minute resolution. Any data points outside ± 3 standard deviations were set to the median value of the prior hour, and any points showing near instantaneous change, as defined by local abs(derivative) > 10^5^ as an arbitrary cutoff, were also set to the median value of the previous hour. Small data interrupts resulting from intermittent data pulls (<10 minutes) were linearly interpolated. Continuous data from p26 to p74 were divided into three equal-length phases: pre to mid puberty (p26 to p41), mid to late puberty (p42 to p58), and late puberty to early adulthood (p59 to p74).

### Wavelet Analyses and Statistics of CBT Data

Briefly, Wavelet Transformation (WT) was used to generate a power estimate, representing amplitude and stability of oscillation at a given periodicity, within a signal at each moment in time. Whereas Fourier transforms allow transformation of a signal into frequency space without temporal position (i.e., using sine wave components of infinite length), wavelets are constructed with amplitude diminishing to 0 in both directions from center. This property permits frequency strength calculation at a given position. In the present analyses we use a Morse wavelet with a low number of oscillations (defined by *β=5* and *γ=3*, the frequencies of the two waves superimposed to create the wavelet(Lilly and Olhede, 2012)), similar to wavelets used in many circadian and ultradian applications(Grant et al., 2020; Leise, 2015, 2013; Lilly and Olhede, 2012; Smarr et al., 2017, 2016). Additional values of *β* (3–8) and *γ* (2–5) did not alter the findings. As WTs exhibit artifacts at the edges of the data being transformed, only the WT of the second through the second to last days of data were analyzed further, from p26 to p74. Periods of 1 to 39 h were assessed. For quantification of spectral differences, WT spectra were isolated in bands; circadian periodicity power was defined as the max power per minute within the 23 to 25 h band, and ultradian periodicity power was defined as the max power per minute in the 1 to 3 h band. The latter band was chosen because this band corresponded with the daily ultradian peak power observed in ultradian rhythms across physiological systems in rats(de Kloet and Sarabdjitsingh, 2008; Grant et al., 2018; Kottler et al., 1989; Sanchez-Alavez et al., 2010).

For statistical comparisons of any two groups, Mann Whitney U (MW) rank sum tests were used to avoid assumptions of normality for any distribution. Non-parametric Kruskal-Wallis tests were used instead of ANOVAs for the same reason; for all Kruskal-Wallis tests, *χ*^2^ and *p* values are listed in the text. All relevant comparisons have the same *n*/group, and thus the same degrees of freedom. Mann Kendall (MK) tests were used to assess trends over time in wavelet power (**Figure 2**) and linear CBT (**Figure 4**) over three equally sized temporal windows, described above. For short term (< 3 days of data) statistical comparisons, 1 data point per 4 hours was used (approximately once per ultradian cycle); for longer term (>3 days of data) statistical comparisons, 1 data point per day was used. Dunn’s test was used for multiple comparisons, and Friedman’s tests were utilized in cases of multiple measurements per individual. Circadian power, visualized in **Figure 2A-D** was smoothed with a 24 h window using the MATLAB function “movmean”. Violin plots, which are similar to box plots with probability density of finding different values represented by width(Violin Plots 101, 2021), were calculated using the MATLAB function “violin” and used to visualize both circadian power (**Figure 2J**) and linear CBT (**Figure 4E-G**). Median daily circadian power regressed against each day’s fE2 for each individual using a mixed effects linear regression (MATLAB function “fitlme”). Individuals were treated as random effects, and fE2/fT and median daily CR power treated as fixed effects (**Figure 2G-H**).

### Estradiol and Testosterone Analysis and Statistics

Fecal estradiol and testosterone concentrations by day of life were averaged across animals by group and plotted with shaded mean ± S.E.M (**Figure 1A,C**). Additionally, in females, data were plotted using a 4-day window for each cycle of life over which fecal samples were collected. As individual estrous cycles are not all aligned in time, samples were assessed in 4-day blocks with the highest value in a collection period (e.g., mid puberty) falling on the third day displayed (**Figure 1B**). That is, during each block, fE2 rose over 3 subsequent days with a decrease on the fourth. For example, if animal 1 began puberty on p30 and exhibited a 4-day window peak of fE2 on p33, then that animal’s “first cycle” would be displayed and averaged into a group representation of first cycle as p31, p32, p33, p34. This strategy enabled group assessment of a pre-pubertal 4-day window, as well as an early, mid, and late pubertal cycle, and an early adulthood cycle for Intact and Intact + C animals.

**Figure 1.**
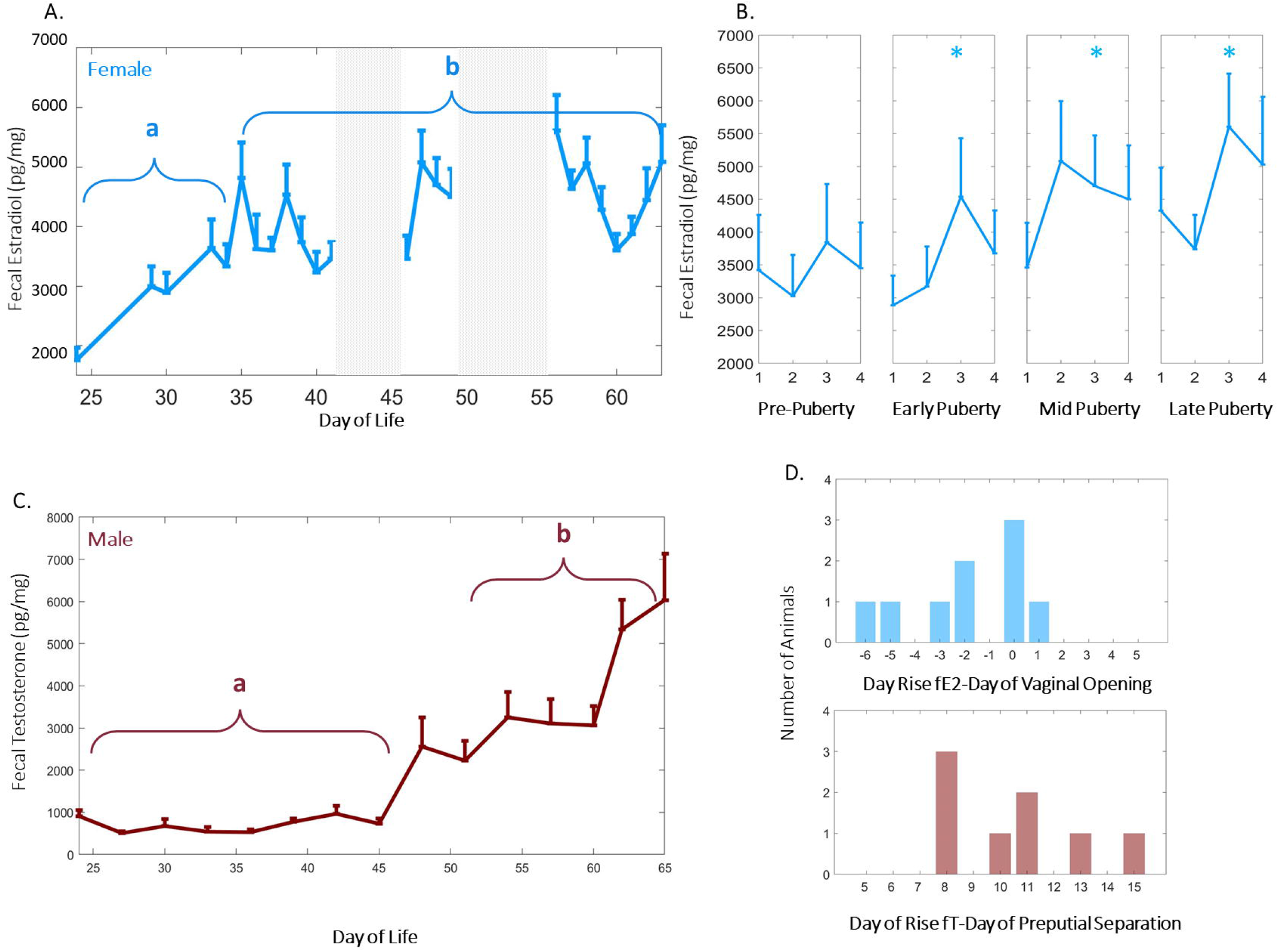
High Frequency Measurement of Fecal Estradiol and Testosterone Enables Monitoring of Estrous Cycle Emergence and Pubertal Progression in Naturalistic Conditions. Group mean (± S.E.M.) of female fecal estradiol (fE2) (blue, A) and male (red, C) fecal testosterone. Fecal testosterone (fT) by day of life differed significantly after p45 (C), whereas the commencement of the ovulatory cycle contributed to variability of female fecal estradiol (A,B). * Letters signify Kruskal Wallis group differences of fT values over the bracketed time region. Group mean and S.E.M. of female (light blue, B) illustrate that fE2 adopts a four-day cycle that stabilizes from early to late puberty, with levels elevated significantly (*p*=0.001) by late adolescence. * Indicates significantly elevated fE2 levels in cycle as compared to pre-pubertal state. FE2 rose 2 standard deviations prior to vaginal opening in most females (D, top), but fT rose 2 standard deviations ~1 to 2 weeks after preputial separation in males (D, bottom).

Day of fE2 or fT rise was defined as the first day fE2 or fT concentration rose > 2 standard deviations above its starting prepubertal value. Relationship to vaginal opening, and preputial separation, are described in **Figure 1D** and **S1**. Group differences in fE2 area under the curve by cycle were assessed using the MATLAB function “trapz” and Kruskal Wallis (KW) tests with Dunn’s *post hoc* correction. Hormone differences by day of life were assessed using Friedman’s test. In order to further assess commencement and stability of estrous cycling after first rise in fE2, metrics were divided into 4 day blocks, with each day labelled 1,2,3, and 4: repeating for subsequent cycle lengths. Groups for statistical comparison were constructed from all data corresponding to 1’s, 2’s, 3’s and 4’s. Friedman’s tests with Dunn’s corrections were used to determine if values associated with each day of cycle (e.g., all day 1’s) varied statistically from other days of the cycle by group.

## Results

### High Frequency Fecal Estradiol and Testosterone Enable Monitoring of Pubertal Progression Under Naturalistic Conditions

In females, fecal estradiol (fE2) increased after p35 (χ^2^= 9.80, *p*=0.001), and exhibited periodic days exhibited elevated fE2 thereafter (*p*=0.03 for days 3 versus day 1 after pubertal onset; **Figure 1A,C-F**). In males, fT increased after p45 (χ^2^= 9.60,*p*=0.002; **Figure 1B**). The relationship between canonical external signs of pubertal onset and fE2/ fT rise was dependent on sex: fE2 rose 2 standard deviations prior to vaginal opening in most females (**Figure 1D**, top), whereas fT rose 2 standard deviations 1 to 2 weeks after preputial separation in males (**Figure 1D**, bottom). Weight trajectories for males and females were typical (**S2**).

### Sex Differences in Circadian Power are Present from Pre-Adolescence through Adulthood

CR, but not UR power rose across early adolescence in both sexes (CR power upward trend *p*=0.009, 0.0012 for females and males, respectively; UR power *p*>0.05 for both sexes; **Figure 2A-C**). Males maintained statistically significantly higher CR power from pre adolescence to mid adolescence and in early adulthood (*χ^2^*= 8.00, 3.78, 16.53; *p*=0.005, 0.052, 4.79*10^-5^ for pre to mid adolescence, mid to late adolescence, and late adolescence to early adulthood, respectively; **Figure 2D-F**). CR power exhibited a non-significant trend toward a 4-day periodic depression after pubertal onset in females (*p*=0.050; **Figure 2E**). CR power was positively correlated with fE2 in adolescent females (*p*=0.04, *r^2^*=0.08, *AIC* = −264; **Figure 2G**), whereas adolescent males exhibited a trend toward a negative correlation between CR power and fE2 (*p*=0.07, *r^2^*=0.09, *AIC*= −132; **Figure 2H**). This pattern was not present prior to pubertal onset, defined by vaginal opening or preputial separation, in either sex (*p*=0.54, *p*=0.89 for females and males, respectively; **Figure 2G-H, insets**).

**Figure 2.**
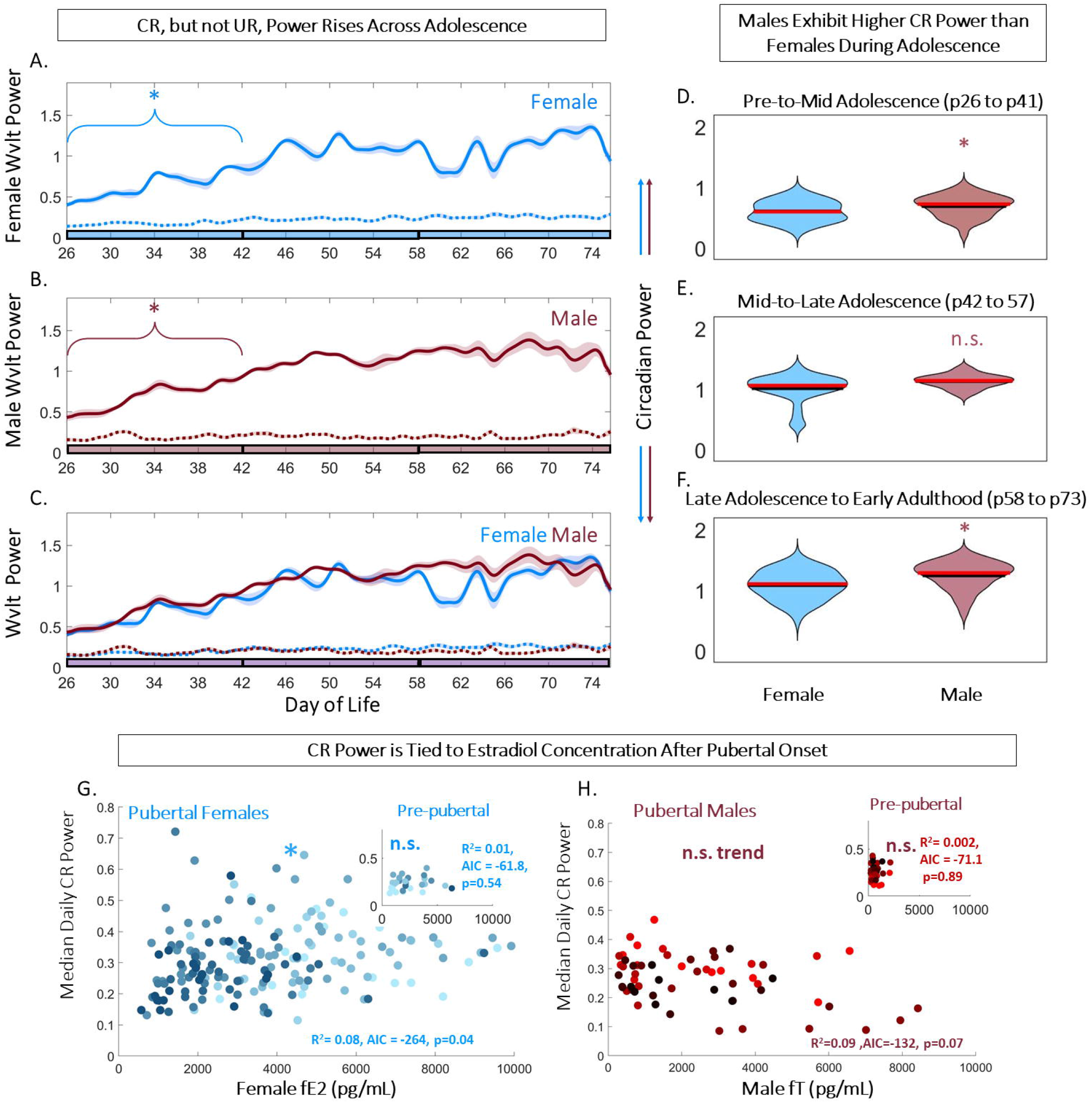
Adolescence Exaggerates Sex Differences in Circadian Power and its Correlation to Sex Steroids. Circadian (CR), but not ultradian (UR) power rises across early adolescence in both sexes (A-C). Linear plots of group mean (± S.E.M.) of CBT CR (solid) and UR (dashed) power in females (blue, A), and males (red, B), overlaid in 2C. * Indicates significant trend over time for the bracketed time (*p*=0.009, 0.0012 for females and males, respectively). Phase of adolescence cutoffs (early to mid, mid to late, and late to adult) are indicated by breaks in the colored x-axis at p42 and p58. Violin plots of CR power illustrate that males maintain significantly (letter indicates group difference) higher CR power than females from early in life (D-F) (*χ^2^*= 8.00, 3.78, 16.53; *p*=0.005, 0.052, 4.79*10^-5^ for pre to mid adolescence, mid to late adolescence, and late adolescence to early adulthood, respectively). Scatters plots of fecal estradiol (fE2) (G) and fecale testosterone (fT) (H) by median daily CR power in females and males, respectively, illustrate a female-specific positive correlation (*p*=0.04, *r^2^*=0.08, *AIC*= −264). This correlation is not present prior to pubertal onset (G,H, insets).

### CBT and Ultradian Power Exhibited Sex-Specific Changes

CBT exhibited an approximately 4-day periodic fluctuation in females, but not males, commencing with the rise in fE2 and vaginal opening (*χ^2^*=11.5, 1.3, *p*=0.003 and *p*>0.05 for females and males, respectively; **Figure 3A-B**). UR power exhibited a comparable 4-day pattern in females (*χ^2^*=8.75, 3.25, *p*=0.005) but not males (*p*>0.05) (**Figure 3A-B**). An FFT of male and female CBT and UR power corroborated these observations; females exhibited statistically greater A.U.C. for 4 to 5 day periodicity of CBT modulation (*χ^2^*=11.29, *p*=8*10^-4^ for sex difference in A.U.C. of 4 to 5 day temperature FFT; **Figure 3C**) and UR modulation (*χ^2^*=9.28, *p*=0.002 for sex difference in A.U.C. of 4 to 5 day UR Power FFT; UR alignment shown in **Figure 3D**). Additionally, females exhibited a statistically significant upward trend in temperature from pre to mid adolescence (*p*=1*10^-5^ to *p*=0.02; mean *p*=0.004), and a significant downward trend in body temperature from mid to late adolescence (*p*=0.019) (**Figure 4A, 4C**). Conversely, males did not exhibit a statistical trend in temperature from early to mid (*p*=0.07) or from mid to late adolescence (*p*=0.12) (**Figure 4B-C**). Violin plots of temperatures across adolescence indicated that females exhibited elevated temperatures compared to males for the entire period of study (*χ^2^*= 25.37, 33.84, 25.52; *p*=9.75*10^-7^, 5.97*10^-9^, 4.37*10^-7^ for pre to mid adolescence, mid to late adolescence, and late adolescence to early adulthood, respectively; **Figure 3A-B, Figure 4D-F**).

**Figure 3.**
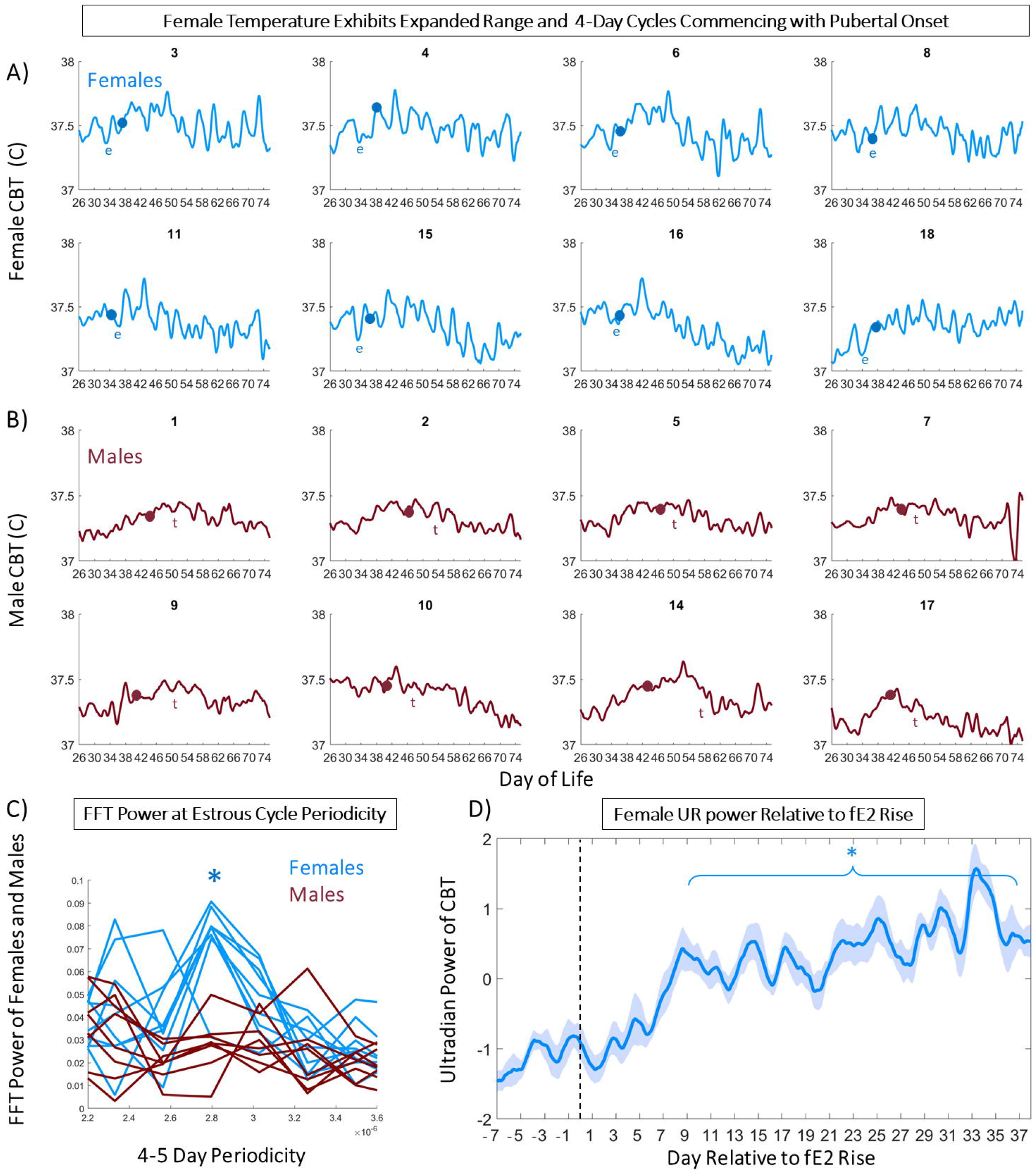
4-Day Patterns of Temperature and Ultradian Power Track Ovulatory Cycles of After Pubertal Onset. Linear plots of smoothed temperature illustrate estrous cycles which onset in time with markers of puberty, vaginal opening and rise in fecal estradiol (fE2) in all individual females (A); and preputial separation and rise in fecal testosterone (fT) in males (B). Dots indicate day of vaginal opening (blue) or preputial separation (red). Letter “e” indicates day of first rise in fE2> 2 standard deviations above the mean, whereas letter “t” indicates day of first rise in fT> 2 standard deviations above the mean. Fast Fourier Transform (FFT) of temperatures of females and males centered on 4 to 5 day periodicities indicate females (blue) exhibit a significant peak compared to males (red) (C). Mean (± S.E.M.) CBT ultradian (UR) power aligned among individuals with reference to first fE2 exhibits onset of 4 to 5 day modulation among females approximately 1 week after fE2 rise (D).

**Figure 4.**
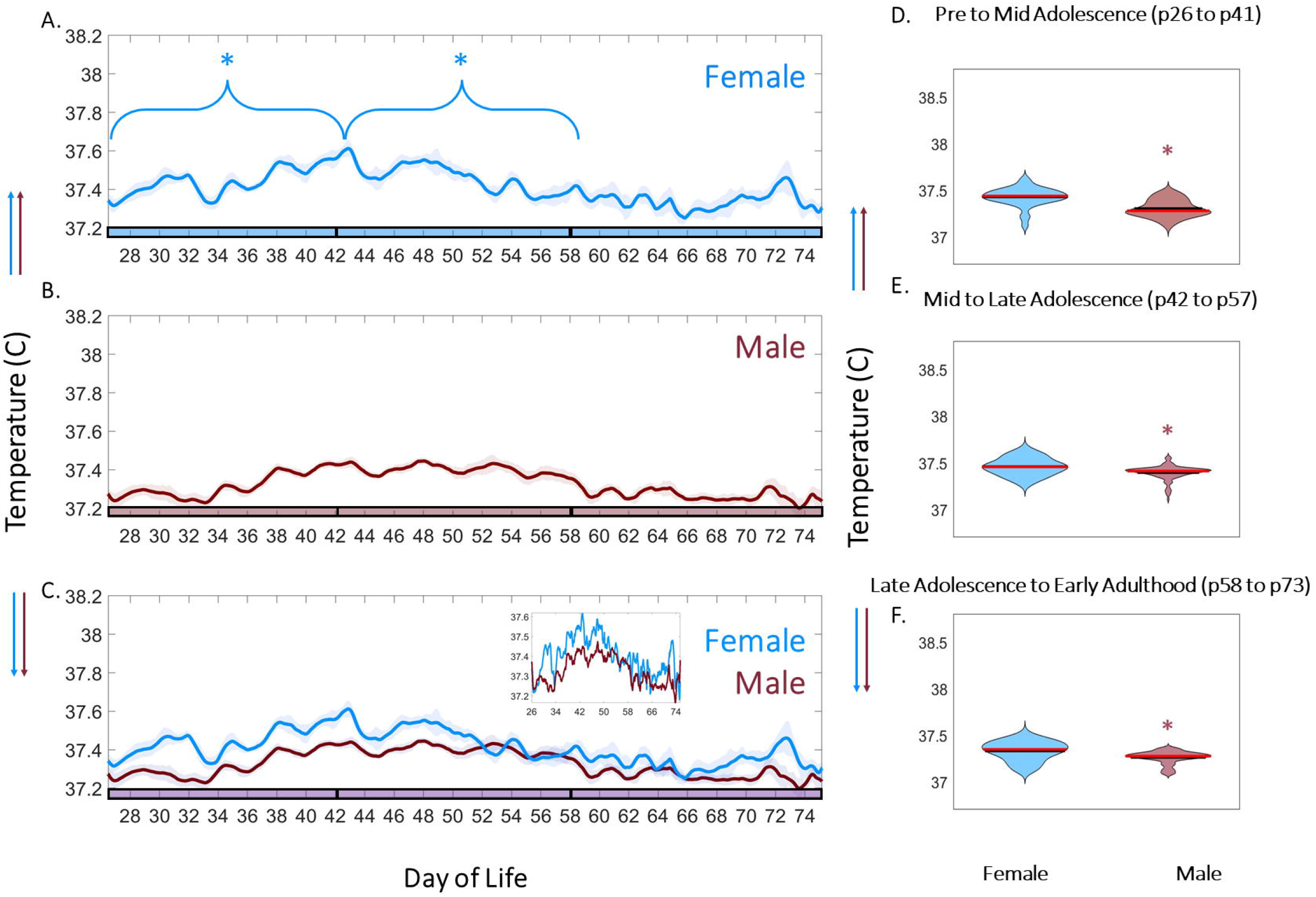
Adolescence is Associated with Sex-Dependent CBT Trends and Levels. CBT Linear group means (± S.E.M.) in females (blue, A), and males (red, B). * Indicates statistically significant MK trend during the bracketed time period. Phase of adolescence cutoffs (pre to mid, mid to late, and late to adult) are indicated by breaks in the colored x-axis at p42 and p58. Females exhibit a statistically significant upward trend in body temperature from pre to mid adolescence (A-C) (*p*=0.004), and a downward trend in body temperature from mid to late adolescence (A) (*p*=0.019). Violin plots of female and male temperatures across adolescence indicate that females exhibit wider ranging (* in D-F indicates significant difference) and elevated temperatures compared to males (C, zoomed inset; D-F) (*χ^2^*= 25.37, 33.84, 25.52; *p*=9.75*10^-7^, 5.97*10^-9^, 4.37*10^-7^ for pre to mid adolescence, mid to late adolescence, and late adolescence to early adulthood, respectively).

## Discussion and Conclusions

The present findings reveal that CBT features gathered in a naturalistic environment can be used to monitor adolescence, despite the additional variability in sex steroid concentrations and environmental factors compared to a traditional laboratory environment. Adolescent trends in CBT and CBT rhythmicity observed in the present study were akin to those of females examined in the laboratory(Grant et al., 2021) with notable sex differences. Males and females exhibited differential trends in and amplitudes of CR and UR power, with the most notable being the rapid onset of females’ 4-day estrous cycle patterning in CBT URs following the rise in fE2 and vaginal opening. CR power increased from pre to mid puberty in both sexes, with females exhibiting higher CBT and lower CR power than males. Despite the observation of higher and more variable fE2 compared to lab-reported values, females retained a statistically significant correlation between fE2 and CR power after pubertal onset(Grant et al., 2021). In contrast to the coordinated patterns in fE2 and CBT in females, coordinated changes in CBT structure and fT were not observed in males. Together, these findings affirm that CBT and CBT rhythmicity remain informative in variable environments, particularly in females, and support the potential for CBT-based monitoring outside of the laboratory environment and across species.

The similarity among the trajectories of circadian power in males and females is intriguing given that fT rose much later in males than fE2 in females(Hagenauer et al., 2011; MacKinnon et al., 1978; Sengupta, 2013). Because the rise in fT was temporally decoupled from rhythmic metrics and preputial separation, a sex-steroid-independent physiological change might drive an early rise CBT rhythmicity (e.g., melatonin(Cavallo, 1993; Rivest et al., 1986) or growth hormone(Dunger et al., 1991; Grant et al., 2018)). Despite remarkable similarities in adolescent circadian, ultradian, and CBT trajectories between the sexes, the presence of elevated CBT (which persists into adulthood(Zuloaga et al., 2009)) and reduced circadian power in females, suggests that continuous-temperature-based diagnostic algorithms should take sex into account during training and validation.

If the features described here have analogous counterparts in human populations, as has recently been shown for continuous temperature for female LH surge anticipation(Grant et al., 2020; Webster and Smarr, 2020), pregnancy(Grant et al., 2021), and fever(Smarr et al., 2020); then this approach can be applied to develop powerful tools to further understand key developmental events. At present, children in developed nations begin puberty at an earlier age than in past decades, attributed to body fat and stress-related factors(Bellis et al., 2006, p. 12; Chittwar et al., 2012; Delemarre-van de Waal et al., 2002; Herbison, 2016; Parent et al., 2003). Additionally, these children are subject to widely varying temporal disruptions in the form of light at night(Casper and Gladanac, 2014; Jain Gupta and Khare, 2020; Smarr and Schirmer, 2018), late meals(Jain Gupta and Khare, 2020), and female hormonal contraceptives(Apter, 2018). Despite the need for monitoring the effects and interactions of these variables on pubertal health, clinicians are equipped with relatively low temporal resolution tools for pubertal staging and diagnosis(Elchuri and Momen, 2020; Klein et al., 2017; Lauffer et al., 2020). Furthermore, the importance of rhythmic stability throughout adolescent development is often not considered by families or pediatricians(Owens and Weiss, 2017).

Peripheral measurements of temperature, such as those from the iButton(Hasselberg et al., 2013) or Oura Ring(Grant et al., 2020; Maijala et al., 2019), could be sufficient for peripubertal detection of temperature and ultradian power rises(Grant et al., 2020), and could be used to develop a population-wide database characterizing features associated with pubertal onset and development. Indeed, rhythmic features of body temperature have already formed the basis of methods for monitoring reproductive health, including pubertal onset(Grant et al., 2021) and contraceptive use in a laboratory setting(Grant et al., 2021), adult fertility in controlled and real world conditions(Grant et al., 2020; Prendergast et al., 2012; Sanchez-Alavez et al., 2011; Smarr et al., 2017), and pregnancy in the laboratory and in small, retrospective cohorts(Grant et al., 2021.; Smarr et al., 2016; Wang et al., 2014). Such tools could be informative and empowering to young people during puberty, potentially anticipating first onset of menses(Fowler et al., 2020; Wartella et al., 2016), impending growth spurts, or for identifying adverse reactions to disruptive behavior(Asimes et al., 2018; Logan et al., 2018) and medication(Apter, 2018). If adopted and studied in teen populations, these metrics could be used to generate the first high-temporal-resolution images of healthy adolescent development and to aid early diagnosis via detection of deviations from a personalized healthy trajectory.

Together, non-invasive sex steroid measurement and chronic observation of CBT rhythms and amplitude represent promising metrics for the detection of pubertal onset and monitoring of the developmental trajectory in both sexes under naturalistic conditions, particularly in females. Future work is needed to determine the extent to which such features are extant and coordinated with markers of puberty in humans, but the present findings in rats suggest the feasibility of such an approach.

## Acknowledgements

The authors would like to thank Andrew Ahn, Ronald Dahl, Frédéric Theunissen, and Albert Qü for their helpful feedback on methods. This work was supported by a Miller Professorship from the Miller Institute for Basic Research at UC Berkeley (LW), and by NIH Grant HD-050470 (LJK).

## Supplemental Figures

**S1:**
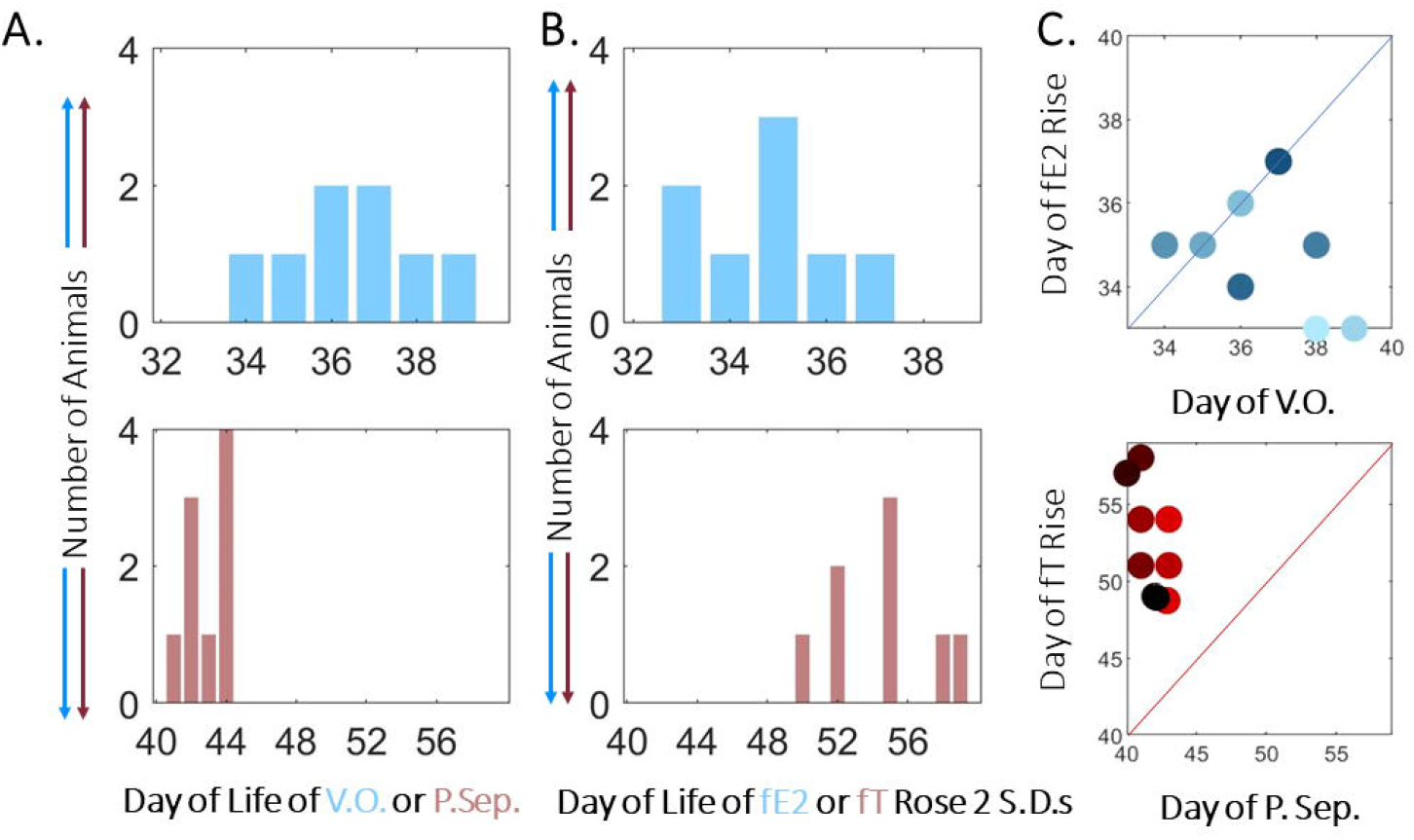
External Pubertal Signs Differ in Time from Sex Steroid Rise. Distribution of the day of vaginal opening (top) or preputial separation (bottom) (A) versus rise in fE2 (top) and rise in fT (bottom) (B) illustrate that females’ external and hormonal pubertal onset markers are more tightly coupled than in males. Scatter plot (C) of day of external marker versus day of hormonal rise >2 standard deviations above the mean.

**S2:**
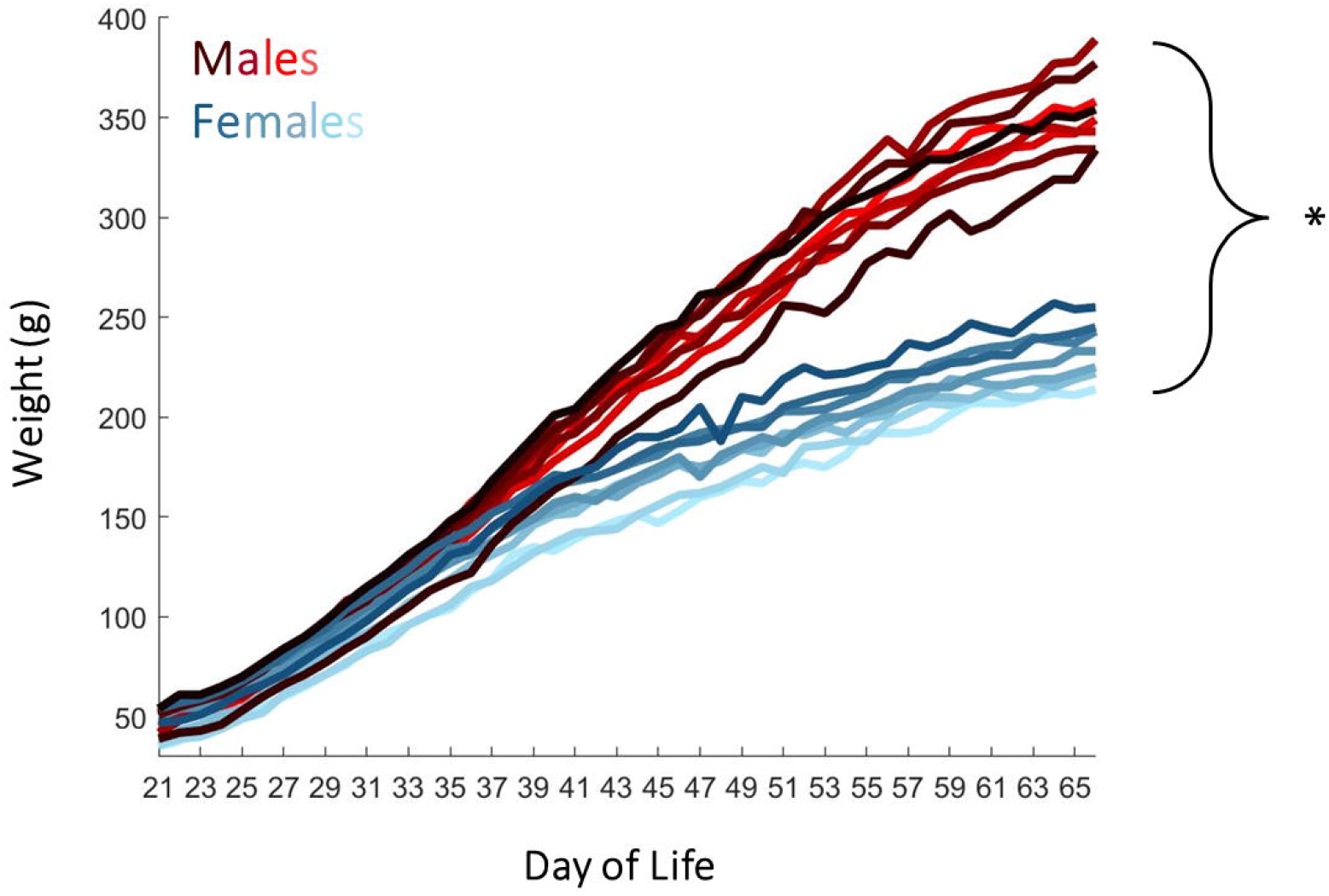
Male and Female Pubertal Weight Trajectory. Plot of day of life and weight for females (blue gradient) and males (red gradient). Male and female weights differ statistically by the approximate time of female pubertal onset, p39 (*p*=0.001).

## References

Akin A, Elstein M (1975) The value of the basal temperature chart in the management of infertility. Int J Fertil 20:122–124.

Albertsson-Wikland K, Rosberg S, Lannering B, Dunkel L, Selstam G, Norjavaara E (1997) Twenty-Four-Hour Profiles of Luteinizing Hormone, Follicle-Stimulating Hormone, Testosterone, and Estradiol Levels: A Semilongitudinal Study throughout Puberty in Healthy Boys. J Clin Endocrinol Metab 82:541–549.

Ankarberg C, Norjavaara E (1999) Diurnal rhythm of testosterone secretion before and throughout puberty in healthy girls: correlation with 17beta-estradiol and dehydroepiandrosterone sulfate. J Clin Endocrinol Metab 84:975–984.

Apter D (2018) Contraception options: Aspects unique to adolescent and young adult. Best Pract Res Clin Obstet Gynaecol 48:115–127.

Asimes A, Kim CK, Cuarenta A, Auger AP, Pak TR (2018) Binge Drinking and Intergenerational Implications: Parental Preconception Alcohol Impacts Offspring Development in Rats. J Endocr Soc 2:672–686.

Auer KE, Kußmaul M, Möstl E, Hohlbaum K, Rülicke T, Palme R (2020) Measurement of Fecal Testosterone Metabolites in Mice: Replacement of Invasive Techniques. Anim Open Access J MDPI 10. azuredominique (2021) azuredominique/Rat-Puberty-Lab-Conditions.

Bakshi VP, Geyer MA (1999) Ontogeny of isolation rearing-induced deficits in sensorimotor gating in rats. Physiol Behav 67:385–392.

Bellis MA, Downing J, Ashton JR (2006) Adults at 12? Trends in puberty and their public health consequences. J Epidemiol Community Health 60:910–911.

Bhavani SV, Carey KA, Gilbert ER, Afshar M, Verhoef PA, Churpek MM (2019) Identifying Novel Sepsis Subphenotypes Using Temperature Trajectories. Am J Respir Crit Care Med 200:327–335.

Boggiano MM, Cavigelli SA, Dorsey JR, Kelley CEP, Ragan CM, Chandler-Laney PC (2008) Effect of a cage divider permitting social stimuli on stress and food intake in rats. Physiol Behav 95:222–228.

Bourguignon JP (1988) Time-related neuroendocrine manifestations of puberty: a combined clinical and experimental approach extracted from the 4th Belgian Endocrine Society lecture. Horm Res 30:224–234.

Brown-Douglas CG, Firth EC, Parkinson TJ, Fennessy PF (2004) Onset of puberty in pasture-raised Thoroughbreds born in southern hemisphere spring and autumn. Equine Vet J 36:499–504.

Buxton CL, Atkinson WB (1948) Hormonal factors involved in the regulation of basal body temperature during the menstrual cycle and pregnancy. J Clin Endocrinol Metab 8:544–549.

Campbell-Page RM, Shaw-Ridley M (2013) Managing Ethical Dilemmas in Community-Based Participatory Research With Vulnerable Populations. Health Promot Pract 14:485–490.

Casper RF, Gladanac B (2014) Introduction: circadian rhythm and its disruption: impact on reproductive function. Fertil Steril 102:319–320.

Cavallo A (1993) Melatonin and human puberty: current perspectives. J Pineal Res 15:115–121.

Chittwar S, Shivprakash, Ammini AC (2012) Precocious puberty in girls. Indian J Endocrinol Metab 16:S188–S191.

Daan S, Slopsema S (1978) Short-term rhythms in foraging behaviour of the common vole, Microtus arvalis. J Comp Physiol 127:215–227.

de Kloet ER, Sarabdjitsingh RA (2008) Everything has rhythm: focus on glucocorticoid pulsatility. Endocrinology 149:3241–3243.

Delemarre-van de Waal HA, van Coeverden SCCM, Engelbregt MTJ (2002) Factors affecting onset of puberty. Horm Res 57 Suppl 2:15–18.

Dunger DB, Matthews DR, Edge JA, Jones J, Preece MA (1991) Evidence for temporal coupling of growth hormone, prolactin, LH and FSH pulsatility overnight during normal puberty. J Endocrinol 130:141–149.

Elchuri SV, Momen JJ (2020) Disorders of Pubertal Onset. Prim Care Clin Off Pract, Adolescent Medicine 47:189–216.

Fowler LR, Gillard C, Morain S (2020) Teenage Use of Smartphone Applications for Menstrual Cycle Tracking. Pediatrics 145.

Franklin AD, Waddell WT, Behrns S, Goodrowe KL (2020) Estrous cyclicity and reproductive success are unaffected by translocation for the formation of new reproductive pairs in captive red wolves (Canis rufus). Zoo Biol 39:230–238.

Garcia J, Rosen G, Mahowald M (2001) Circadian rhythms and circadian rhythm disorders in children and adolescents. Semin Pediatr Neurol 8:229–240.

Gear AJCH (2014) Wearables Are Totally Failing the People Who Need Them Most [WWW Document]. WIRED. URL https://www.wired.com/2014/11/where-fitness-trackers-fail/ (accessed 8.4.17).

Goh GH, Maloney SK, Mark PJ, Blache D (2019) Episodic Ultradian Events—Ultradian Rhythms. Biology 8:15.

Grant AD, Smarr B (2021). Feasibility of Continuous Distal Body Temperature for Passive, Early Pregnancy Detection BioArxiv, 24.

Grant AD, Newman M, Kriegsfeld LJ (2020) Ultradian rhythms in heart rate variability and distal body temperature anticipate onset of the luteinizing hormone surge. Sci Rep 10:20378.

Grant AD, Wilbrecht L, Kriegsfeld LJ (2021) Adolescent Development of Biological Rhythms: Estradiol Dependence and Effects of Combined Contraceptives. bioRxiv 2021.07.20.453145.

Grant AD, Wilsterman K, Smarr BL, Kriegsfeld LJ (2018) Evidence for a Coupled Oscillator Model of Endocrine Ultradian Rhythms. J Biol Rhythms 33:475–496.

Grant AD, Wolf GI, Nebeker C (2019) Approaches to governance of participant-led research: a qualitative case study. BMJ Open 9:e025633.

Hagenauer MH, King AF, Possidente B, McGinnis MY, Lumia AR, Peckham EM, Lee TM (2011) Changes in circadian rhythms during puberty in Rattus norvegicus: developmental time course and gonadal dependency. Horm Behav 60:46–57.

Harper JM, Austad SN (2000) Fecal glucocorticoids: a noninvasive method of measuring adrenal activity in wild and captive rodents. Physiol Biochem Zool PBZ 73:12–22.

Hasselberg MJ, McMahon J, Parker K (2013) The validity, reliability, and utility of the iButton® for measurement of body temperature circadian rhythms in sleep/wake research. Sleep Med 14:5–11.

Herbison AE (2016) Control of puberty onset and fertility by gonadotropin-releasing hormone neurons. Nat Rev Endocrinol 12:452–466.

Hoogenboom I, Daan S, Dallinga JH, Schoenmakers M (1984) Seasonal Change in the Daily Timing of Behaviour of the Common Vole, Microtus arvalis. Oecologia 61:18–31.

Jain Gupta N, Khare A (2020) Disruption in daily eating-fasting and activity-rest cycles in Indian adolescents attending school. PloS One 15:e0227002.

Joyce DS, Zele AJ, Feigl B, Adhikari P (2020) The accuracy of artificial and natural light measurements by actigraphs. J Sleep Res 29:e12963.

Kalliokoski O, Teilmann AC, Abelson KSP, Hau J (2015) The distorting effect of varying diets on fecal glucocorticoid measurements as indicators of stress: A cautionary demonstration using laboratory mice. Gen Comp Endocrinol 211:147–153.

Kim HJ, Harrington ME (2008) Neuropeptide Y-deficient mice show altered circadian response to simulated natural photoperiod. Brain Res 1246:96–100.

Klein DA, Emerick JE, Sylvester JE, Vogt KS (2017) Disorders of Puberty: An Approach to Diagnosis and Management. Am Fam Physician 96:590–599.

Kottler ML, Coussieu C, Valensi P, Levi F, Degrelle H (1989) Ultradian, circadian and seasonal variations of plasma progesterone and LH concentrations during the luteal phase. Chronobiol Int 6:267–277.

Kriegsfeld-Lab - Overview [WWW Document] (n.d.) . GitHub. URL https://github.com/Kriegsfeld-Lab (accessed 5.24.21).

Lafaille M, Gouat P, Féron C (2015) Efficiency of delayed reproduction in Mus spicilegus. Reprod Fertil Dev 27:491–496.

Lauffer P, Mooij CF, Zwaveling-Soonawala N, Trotsenburg ASP van (2020) Reforming the male Tanner genital scale. J Pediatr Endocrinol Metab 33:425–426.

Leise TL (2015) Chapter Five - Wavelet-Based Analysis of Circadian Behavioral Rhythms In: Methods in Enzymology, Circadian Rhythms and Biological Clocks, Part A (Sehgal A ed), pp95–119. Academic Press.

Leise TL (2013) Wavelet analysis of circadian and ultradian behavioral rhythms. J Circadian Rhythms 11:5.

Lewis R, Curtis JT (2016) Male prairie voles display cardiovascular dipping associated with an ultradian activity cycle. Physiol Behav 156:106–116.

Lilly JM, Olhede SC (2012) Generalized Morse Wavelets as a Superfamily of Analytic Wavelets. IEEE Trans Signal Process 60:6036–6041.

Logan RW, Hasler BP, Forbes EE, Franzen PL, Torregrossa MM, Huang YH, Buysse DJ, Clark DB, McClung CA (2018) Impact of Sleep and Circadian Rhythms on Addiction Vulnerability in Adolescents. Biol Psychiatry 83:987–996.

Lv X et al. (2020) Reprogramming of ovarian granulosa cells by YAP1 leads to development of high-grade cancer with mesenchymal lineage and serous features. Sci Bull 65:1281–1296.

MacKinnon PCB, Puig-Duran E, Laynes R (1978) Reflections on the attainment of puberty in the rat: have circadian signals a role to play in its onset? Reproduction 52:401–412.

Maijala A, Kinnunen H, Koskimäki H, Jämsä T, Kangas M (2019) Nocturnal finger skin temperature in menstrual cycle tracking: ambulatory pilot study using a wearable Oura ring. BMC Womens Health 19:150.

Mathew L, Gaikwad A, Gonzalez A, Nugent EK, Smith JA (2017) Evaluation of Active Hexose Correlated Compound (AHCC) in Combination With Anticancer Hormones in Orthotopic Breast Cancer Models. Integr Cancer Ther 16:300–307.

Meijer JH, Michel S, Vanderleest HT, Rohling JHT (2010) Daily and seasonal adaptation of the circadian clock requires plasticity of the SCN neuronal network. Eur J Neurosci 32:2143–2151.

Millspaugh JJ, Washburn BE (2003) Within-sample variation of fecal glucocorticoid measurements. Gen Comp Endocrinol 132:21–26.

Mohawk JA, Green CB, Takahashi JS (2012) Central and peripheral circadian clocks in mammals. Annu Rev Neurosci 35:445–462.

Neurodevelopmental Consequences of Maternal Omega-3 Fatty Acid Deficiency - ProQuest [WWW Document] (n.d.). URL https://search.proquest.com/openview/2b9eaffc7b3f492b60daf6e1fababebd/1?pq-origsite=gscholar&cbl=2026366&diss=y (accessed 3.3.21).

Norjavaara E, Ankarberg C, Albertsson-Wikland K (1996) Diurnal rhythm of 17 beta-estradiol secretion throughout pubertal development in healthy girls: evaluation by a sensitive radioimmunoassay. J Clin Endocrinol Metab 81:4095–4102.

Owens JA, Weiss MR (2017) Insufficient sleep in adolescents: causes and consequences. Minerva Pediatr 69:326–336.

Parent A-S, Teilmann G, Juul A, Skakkebaek NE, Toppari J, Bourguignon J-P (2003) The Timing of Normal Puberty and the Age Limits of Sexual Precocity: Variations around the World, Secular Trends, and Changes after Migration. Endocr Rev 24:668–693.

Park JH, Spencer EM, Place NJ, Jordan CL, Zucker I (2003) Seasonal control of penile development of Siberian hamsters (Phodopus sungorus) by daylength and testicular hormones. Reprod Camb Engl 125:397–407.

Steroid Solid Extraction [WWW Document]. Arbor Assays. URL https://www.arborassays.com/resource/steroid-solid-extraction/ (accessed 3.3.21).

Prendergast BJ, Beery AK, Paul MJ, Zucker I (2012) Enhancement and Suppression of Ultradian and Circadian Rhythms across the Female Hamster Reproductive Cycle. J Biol Rhythms 27:246–256.

Righetti F, Tybur J, Van Lange P, Echelmeyer L, van Esveld S, Kroese J, van Brecht J, Gangestad S (2020) How reproductive hormonal changes affect relationship dynamics for women and men: A 15-day diary study. Biol Psychol 149:107784.

Rivest RW, Aubert ML, Lang U, Sizonenko PC (1986) Puberty in the rat: modulation by melatonin and light. J Neural Transm Suppl 21:81–108.

Rueda-Quijano SM, Amador-Ariza MA, Arboleda AM, Otero J, Cohen D, Camacho PA, Jaramillo PL (2019) [Concordance of the assessment of pubertal development with the Tanner scale between adolescents and a trained physician]. Rev Peru Med Exp Salud Publica 36:408–413.

Sanchez-Alavez M et al. (2010) Insulin causes hyperthermia by direct inhibition of warm-sensitive neurons. Diabetes 59:43–50.

Sanchez-Alavez M, Alboni S, Conti B (2011) Sex- and age-specific differences in core body temperature of C57Bl/6 mice. Age Dordr Neth 33:89–99.

Sengupta P (2013) The Laboratory Rat: Relating Its Age With Human’s. Int J Prev Med 4:624–630.

Shirtcliff EA, Dahl RE, Pollak SD (2009) Pubertal development: correspondence between hormonal and physical development. Child Dev 80:327–337.

Silva C-C, Domínguez R (2020) Clock control of mammalian reproductive cycles: Looking beyond the pre-ovulatory surge of gonadotropins. Rev Endocr Metab Disord 21:149–163.

Smarr BL, Aschbacher K, Fisher SM, Chowdhary A, Dilchert S, Puldon K, Rao A, Hecht FM, Mason AE (2020) Feasibility of continuous fever monitoring using wearable devices. Sci Rep 10.

Smarr BL, Grant AD, Zucker I, Prendergast BJ, Kriegsfeld LJ (2017) Sex differences in variability across timescales in BALB/c mice. Biol Sex Differ 8:7.

Smarr BL, Schirmer AE (2018) 3.4 million real-world learning management system logins reveal the majority of students experience social jet lag correlated with decreased performance. Sci Rep 8:4793.

Smarr BL, Zucker I, Kriegsfeld LJ (2016) Detection of Successful and Unsuccessful Pregnancies in Mice within Hours of Pairing through Frequency Analysis of High Temporal Resolution Core Body Temperature Data. PloS One 11:e0160127.

Steadman C (2019) The effect of spinal cord injury on sexual function. University of Louisville.

Steadman CJ, Hoey RF, Montgomery LR, Hubscher CH (2019) Activity-Based Training Alters Penile Reflex Responses in a Rat Model of Spinal Cord Injury. J Sex Med 16:1143–1154.

Steiger SS, Valcu M, Spoelstra K, Helm B, Wikelski M, Kempenaers B (2013) When the sun never sets: diverse activity rhythms under continuous daylight in free-living arctic-breeding birds. Proc Biol Sci 280:20131016.

Stothard ER, McHill AW, Depner CM, Birks BR, Moehlman TM, Ritchie HK, Guzzetti JR, Chinoy ED, LeBourgeois MK, Axelsson J, Wright Jr. KP (2017) Circadian Entrainment to the Natural Light-Dark Cycle across Seasons and the Weekend. Curr Biol 27:508–513.

The Mammalian Circadian Timing System: Organization and Coordination of Central and Peripheral Clocks (2010). Annu Rev Physiol 72:517–549.

Touma C, Palme R, Sachser N (2004) Analyzing corticosterone metabolites in fecal samples of mice: a noninvasive technique to monitor stress hormones. Horm Behav 45:10–22.

Vidal JD (2017) The Impact of Age on the Female Reproductive System. Toxicol Pathol 45:206–215.

Violin Plots 101: Visualizing Distribution and Probability Density [WWW Document] (n.d.). URL https://mode.com/blog/violin-plot-examples/ (accessed 3.3.21).

Wang ZY, Cable EJ, Zucker I, Prendergast BJ (2014) Pregnancy-induced changes in ultradian rhythms persist in circadian arrhythmic Siberian hamsters. Horm Behav 66:228–237.

Wartella E, Rideout V, Montague H, Beaudoin-Ryan L, Lauricella A (2016) Teens, health and technology: A national survey. Media Commun 4:13–23.

Webster WW, Smarr B (2020) Using Circadian Rhythm Patterns of Continuous Core Body Temperature to Improve Fertility and Pregnancy Planning. J Circadian Rhythms 18:5.

Williams H, Dacks PA, Rance NE (2010) An Improved Method for Recording Tail Skin Temperature in the Rat Reveals Changes During the Estrous Cycle and Effects of Ovarian Steroids. Endocrinology 151:5389–5394.

Woodruff JA, Lacey EA, Bentley G (2010) Contrasting fecal corticosterone metabolite levels in captive and free-living colonial tuco-tucos (Ctenomys sociabilis). J Exp Zool Part Ecol Genet Physiol 313A:498–507.

Woodruff JA, Lacey EA, Bentley GE, Kriegsfeld LJ (2013) Effects of social environment on baseline glucocorticoid levels in a communally breeding rodent, the colonial tuco-tuco (Ctenomys sociabilis). Horm Behav 64:566–572.

Zuloaga DG, McGivern RF, Handa RJ (2009) Organizational influence of the postnatal testosterone surge on the circadian rhythm of core body temperature of adult male rats. Brain Res 1268:68–75.

